# Interindividual vocal tract diversity influences the phonetic diversification of spoken languages

**DOI:** 10.1101/2024.07.26.605254

**Authors:** Steven Moran, Tiena Danner, Marcia Ponce de León, Christoph Zollikofer

## Abstract

Vocal tracts, like other physical traits, exhibit extensive interindividual variation. However, little is known about how this variation translates into phonetic diversity between speakers, and ultimately, across languages. We demonstrate that different vocal tract shapes and associated articulatory strategies leave consistent acoustic signatures, which show congruent patterns of phonetic variation both within and across speech communities. Recalling a central tenet of evolutionary biology - that within-group variation feeds processes of between-group diversification - our findings suggest that the sounds of language evolve according to a neutral-like evolutionary process. Our “neutral-like evolution model” serves as a null hypothesis for disentangling the biological versus cultural mechanisms at play in phonetic diversification, where deviations from the neutral expectation indicate culturally mediated processes of language change at work.

---

TEXTBOOK depictions imply a standard human vocal tract morphology^1–3^. However, there is extensive interindividual variation in terms of the size and shape of the hard and soft tissues involved in speech production^4, 5^. Both “nature” (natural selection, genetic drift)^6–9^ and “nurture” (environment, culture, lifestyle)^10–13^ shape an individual’s vocal tract morphology and imbue their voice with acoustic markers (age, gender, sex, size)^14, 15^ and speaker-specific voice qualities^16, 17^, making them sound unique^18^.

Despite considerable interindividual variation, however, speakers within a speech community reach acoustically similar targets, which they perceive as the same contrastive speech sounds, i.e., phonemes. While speaker communities thus tend to conserve phonetic (as well as other linguistic) properties, it is well known that language properties are constantly changing through time^19, 20^, and a key question to answer is how biology, environment, and culture together bring about these changes.

Here we focus on the biological side of the question, asking specifically how vocal tract variation within a speaker community influences phonetic variation, and how phonetic variation within speaker communities gives rise to variation between communities, and eventually between languages. To the best of our knowledge, these interlinked questions have not yet been addressed comprehensively and empirically. Previous studies have focused on population-level differences in vocal tract shape^13, 18, 21^, on specific areas of the vocal tract^5, 21^, differences related to sex, age and size^14, 22, 23^, have used modeling and simulation-based approaches^21, 24–27^, or are preliminary^28–31^.

In this study, we quantify the origins and propagation of variation across the successive levels of speech production; from variation in vocal tract shape and articulatory gestures to variation in the associated phonetic output within a speaker community, and from community-level to language-level phonetic variation. We use the five cardinal vowels (i,e,a,o,u) as a test case, because they are well characterized in terms of articulation and acoustics, and are shared by a wide variety of typologically diverse languages.

To quantify articulatory-acoustic relationships, we propose a new approach, *morpho-phonetics*, that combines geometric morphometric analysis^32^ of the shape of the vocal tract in real-time Magnetic Resonance Imaging (rtMRI)^33^ with spectral analysis of the concurrent audio signal during speech articulation (Fig. 1, Supplementary Fig. 1). We examined the morpho-phonetic covariation of the cardinal vowels as pronounced by 32 speakers (16 women and 16 men) of American English from different ethnic and linguistic backgrounds. Vowels were extracted from the carrier words (‘beat’, ‘bait’, ‘pot’, ‘boat’, ‘boot’) from the USC 75-Speaker Speech MRI Database^33^.

**Figure 1.**
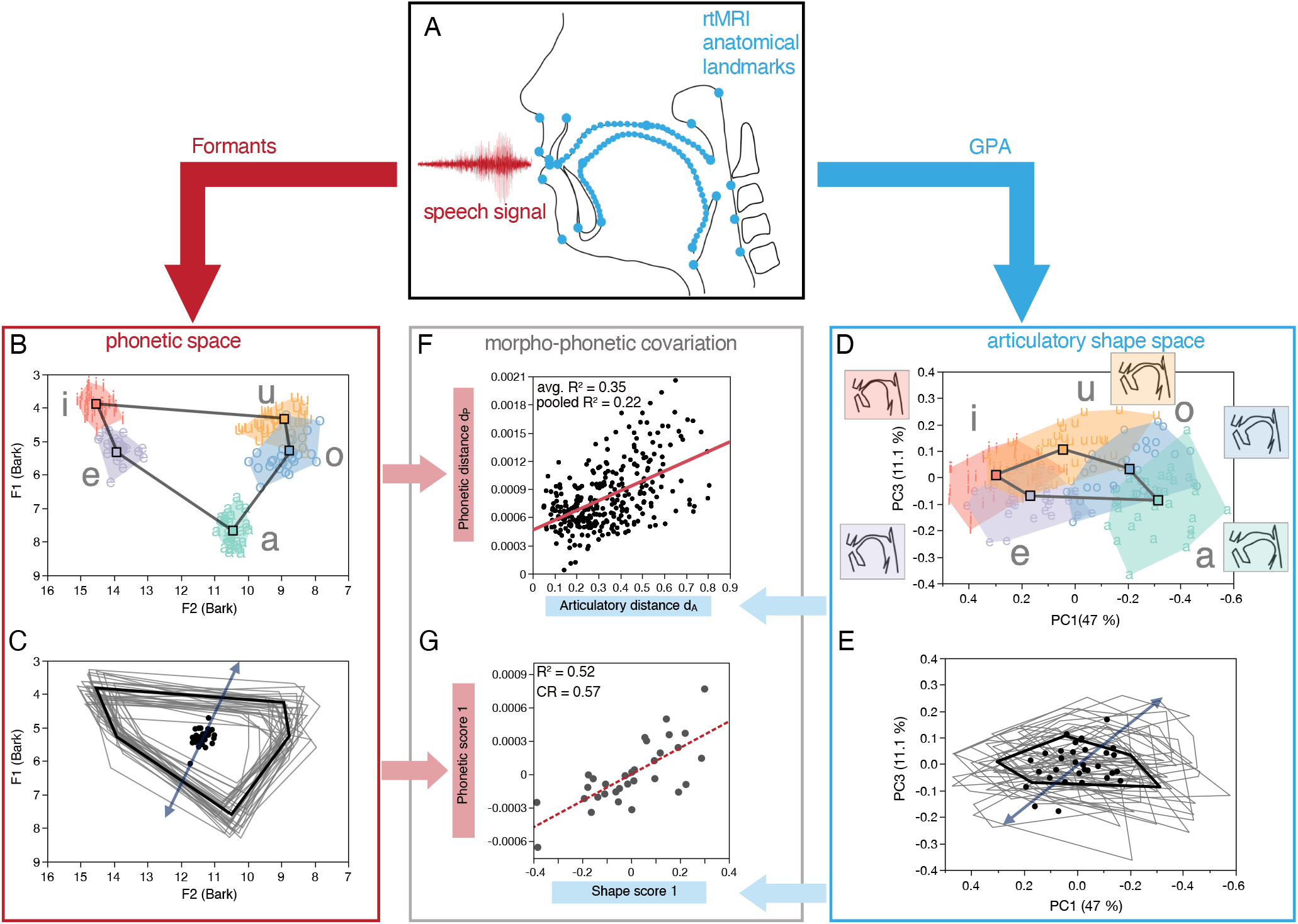
Morpho-phonetic co-analysis of vowel production. **A**: Anatomical landmarks (blue dots) are placed on midsagittal MRI images of the vocal tract recorded during speech production. At each corresponding timestamp, acoustic analysis (formant frequency extraction) is undertaken on denoised audio recordings. **B**-**C**: Resulting acoustic phonetic variation in formant space rendered as interindividual variation of vowels (**B**) and vowel polygons (**C**). Centroids of individual acoustic vowel polygons (black dots; **C**) serve as proxies of acoustic timbre. **D**-**E**: Corresponding variation in articulatory shape space (given by principal components PC1 and PC3) rendered as interindividual variation of vowels (**D**; inset graphs represent mean shape of vocal tract for each vowel) and vowel polygons (**E**). Centroids of individual articulatory vowel polygons (black dots) serve as proxies of the resting-state vocal tract morphologies. In **D** and **B**, note the reduction of within-vowel variation relative to between-vowel variation in phonetic space compared to articulatory shape space, which reflects articulatory adaptation to target vowels. **F**-**G**: Morpho-phonetic covariation. **F**: Linear regressions of phonetic vs. articulatory distances between vowels (individual regressions with average *R*^*2*^ = 0.35; pooled regression with *R*^*2*^ = 0.27 (red line)). **G**: Partial least-squares (PLS) analysis of the individuals’ phonetic vs. articulatory vowel polygon data, reflecting residual morpho-phonetic covariation not compensated by articulatory adaptation. Details in main text and Materials and Methods.

For each vowel spoken by each individual, the articulatory vocal tract shape and its corresponding phonetic output were sampled at synchronous time stamps. Physically, vowel-specific articulation is achieved by coordinated jaw, lip, tongue and velum movements, whose interplay determines the shape of the vocal tract and its acoustic output. Here, vocal tract shape is quantified with a set of anatomical landmarks (*N*=95) (Fig. 1A, Supplementary Fig. 2, Supplementary Table 1) placed on each rtMRI frame, and shape variation is analyzed and visualized with methods of geometric morphometrics (Fig. 1, A, D, E). Articulation results in an acoustic signal (Fig. 1, A, B, C), which is quantified here with vocal tract resonance frequencies, i.e. formants F1-F4, and the fundamental frequency, F0.

Geometric-morphometric analysis of the vocal tract as well as formant analysis yield multidimensional data sets. We use principal components analysis (PCA) as a dimension reduction technique to explore the data structures and to characterize the principal patterns of variation in the data. Covariation between articulatory gestures and phonetic output (i.e., morpho-phonetic covariation) is examined with partial least-squares analysis (PLS). We focus on those patterns of variation that are shared by all individuals, independent of sex, age, and vocal tract size. The workflow and the main results of these analyses are visualized in Fig. 1; details are provided in Materials and Methods. < Insert Figure 1 about here >

## Articulatory and acoustic vowel spaces are similar

Figure 1B shows the acoustic variation given by the first two formants (F1 and F2) in Bark scale^34^. The five vowels are represented by separated data clusters and their centroids form together the well-known acoustic vowel polygon. Figure 1D shows the associated articulatory variation along principal components PC1 and PC3 of shape space (58.1% of total variation). PC2 accounts for interindividual differences unrelated to vowel articulation (13.3% of total variation; see Supplementary Fig. 3A). The articulatory shape space shows partially overlapping vowel data clusters, but notably the centroids of the clusters also form a vowel polygon, which has a similar geometric arrangement as the acoustic vowel polygon. The articulatory and acoustic vowel polygons therefore have corresponding shapes. To quantify the correspondence between articulatory and acoustic vowel polygons, we study how an individual’s articulatory discrimination between vowels translates into acoustic discrimination. To do so, we compute for each individual all ten between-vowel distances in both articulatory and acoustic spaces. Figure 1F shows that articulatory distances correlate with acoustic distances (average of individual speaker correlations: *R*^*2*^ = 0.35; pooled data: *R*^*2*^ = 0.22).

## Articulatory gestures are adapted to speaker-specific vocal tract morphology

The moderate correlation between articulatory and acoustic contrasts (Fig. 1 F) reflects the fact that the individuals in our sample have widely different vocal tract morphologies, and they use a remarkably wide range of articulatory gestures to produce any given target vowel (see Supplementary Figs. 4–8). Therefore, we investigate how individual vocal tract morphology influences speakers’ articulatory gestures, and how these gestures translate into acoustics. A comparison of Figures 1B and 1D shows that there is substantially more articulatory than acoustic variation around corresponding vowel centroids. To quantify this effect, we evaluate the ratio of within-vowel to between-vowel variance (variance ratio; VR). We find that VR is consistently larger in articulatory space (VR_articulatory_ = 0.548) than in acoustic space (VR_formant_ = 0.065; see also Supplementary Table 2). Thus, a wide range of individual articulatory gestures results in a more narrow range of acoustic variation, supporting earlier evidence that individuals adapt their articulatory gestures to produce similar sounding vowels^35, 36^. To further explore the inner workings of articulatory adaptation, we examine how an individual’s palate shape influences the shape of their tongue during vowel production (Fig. 2). To do so, we calculate the mean shape of each individual’s palate and tongue, averaged over all vowels. PLS analysis shows a significant influence of palate shape on tongue shape (Fig. 2A, see Methods). Interindividual differences in palate shape therefore translate into systematic differences in articulatory tongue shape. Specifically, individuals with short, rounded hard palates produce more rounded tongue gestures than individuals with elongated and shallow palates (Fig. 2B). Fig. 2C indicates that articulatory adaptation of the tongue tends to maintain a standard cross-sectional air tube profile in the anterior region of the oral cavity, but less so in the posterior region. Overall, differences in vocal tract morphology are only in part acoustically compensated through articulatory adaptation. < Insert Figure 2 about here >

**Figure 2.**
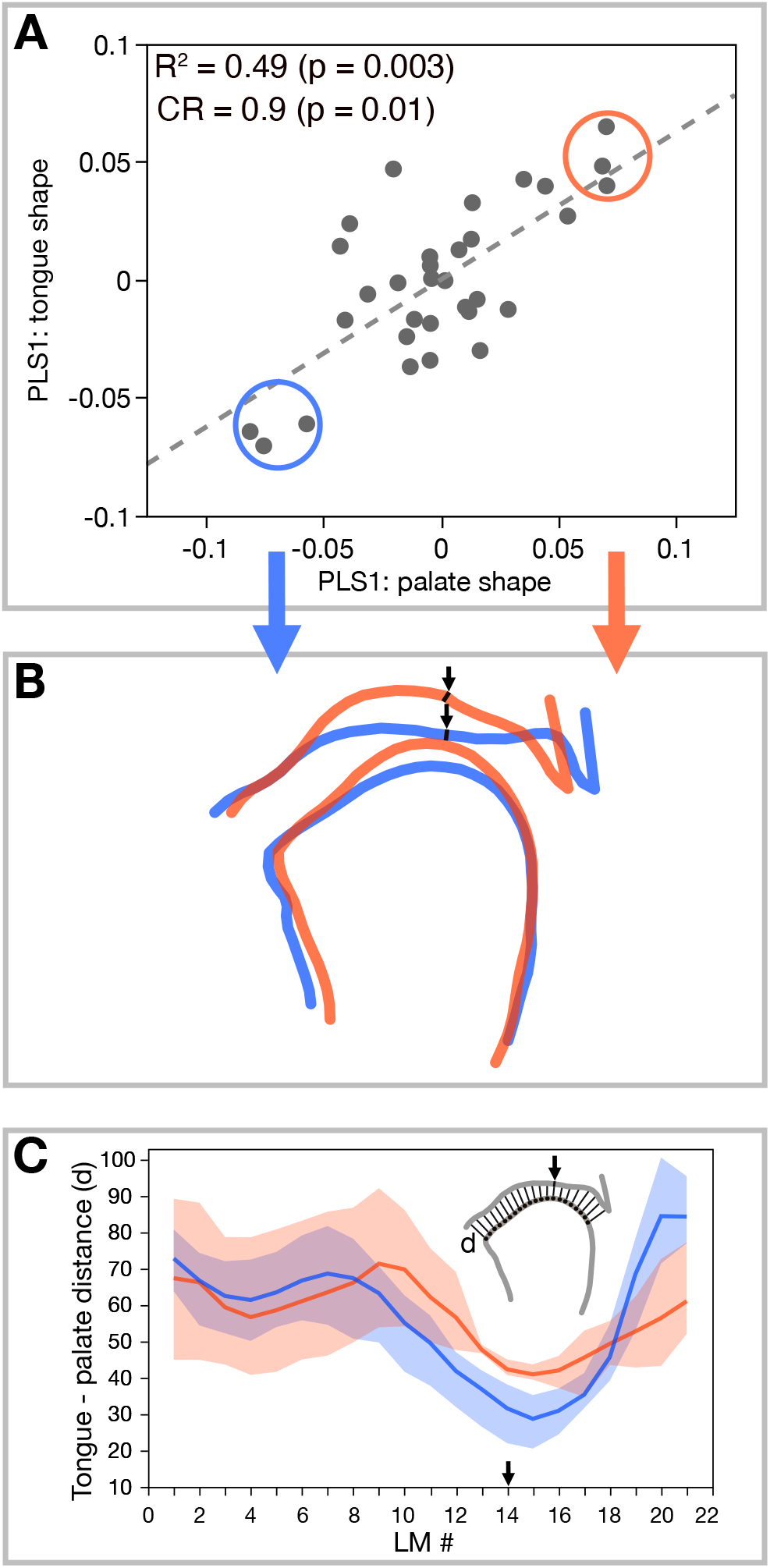
Articulatory adaptation of the tongue to individual palate shape. **A**: Covariation between tongue and palate shapes is computed with partial least squares analysis (PLS). The first pair of PLS components (PLS1) comprises 67.5% of the total covariation. **B**: Articulatory adaptation is visualized for the opposite ends of the morphological variation in **A** (average morphologies of the three individuals comprised in red and blue circles, respectively); black arrows indicate the border between the hard and soft palate. Note that in a short/rounded palate (red) the dorsum of the tongue assumes a more rounded shape. **C**: Air tube cross-sectional profiles derived from the morphologies in **B** (red/blue lines and bands represent group-specific averages and ranges). LM # indicates landmark position along tongue dorsum; **d** denotes tongue-palate distance shown in the inset graph.

## Individual vocal tract morphology results in acoustic differences between speakers

Next we ask how the residual, uncompensated acoustic differences between individuals are related to the underlying differences in their vocal tract morphologies. To do so, we investigate the covariation between individual vowel polygon geometries in articulatory and formant space (Fig. 1G). A lack of covariation would indicate that the acoustic properties of the target vowels are independent of their individual articulatory properties, thus fully compensated for individual vocal tract differences. However, PLS analysis shows significant covariation (Fig. 3A, see Methods), confirming that articulatory adaptation is only partial (Fig. 2B, C), such that individual differences in the vocal tract (Fig. 3A) lead to systematic acoustic differences between speakers (Fig. 3B). This effect is visualized by the real-life morphological and acoustic properties of individuals at opposing ends of the data scatter in Figures 3B and 3C: those with rounded/short versus shallow/elongated palates. Specifically, differences in palate shape result in different timbre of the voice (as indicated by shifts in the position of the vowel polygon in acoustic space), different acoustic range (change in the size of the polygon), and different acoustic contrasts between vowels (change in the shape of the vowel polygon; Fig. 3, B-C). In sum, our analyses show that interindividual variation in vocal tract morphology generates correlated variation in vowel acoustics. < Insert Figure 3 about here >

**Figure 3.**
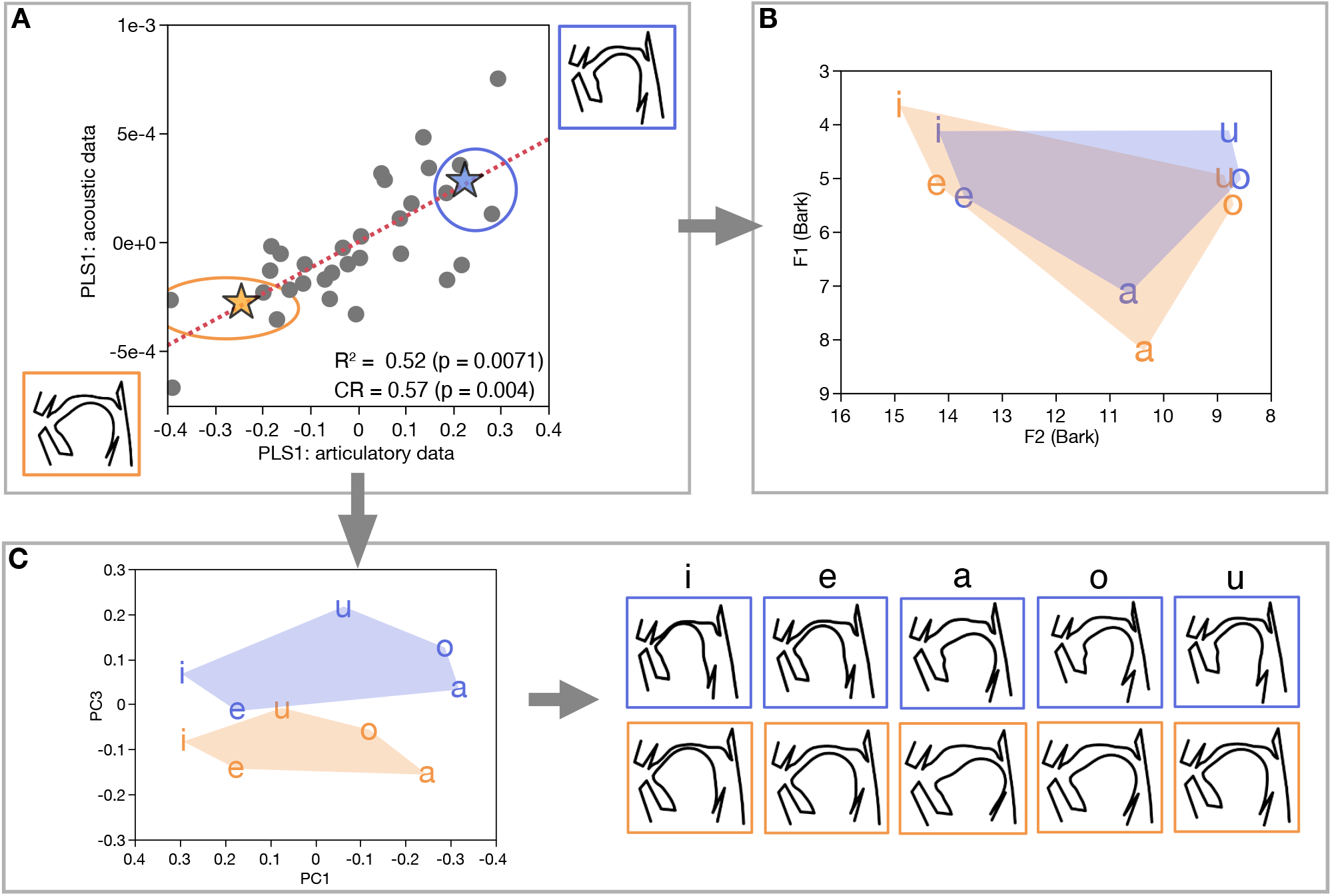
Patterns of uncompensated morpho-phonetic covariation. **A**: PLS analysis of the covariation between articulatory and acoustic vowel polygons (same data as in Fig. 1G). The first pair of PLS components (PLS1) comprises 49.8% of the total covariation. Each data point represents joint articulatory and acoustic data of one individual. Orange/blue stars represent the mean of individuals at the opposite ends of the data scatter (individuals in orange/blue circles) and are used for the visualization of contrasting articulatory morphologies (**C**) and corresponding acoustics (**B**). **B**: Corresponding acoustic vowel polygons in F1-F2 formant space (Bark scale). **C**: Articulatory vowel polygons and associated articulatory differences between individuals with rounded/short and shallow/elongate palates.

## Vocal tract variation influences within- and between-language phonetic diversity

Figure 3 has shown for our morpho-phonetic sample of English speakers (MP-E) that vocal tract variation systematically conditions vowel variation.

Yet the more general question remains: how does vocal tract variation and the ensuing acoustic variation between speakers ultimately influence phonetic diversification across languages? Given the current lack of cross-linguistic morpho-phonetic data, we address this question by combining the MP-E data with the multilingual speech data in the VoxCommunis Corpus (VC-C)^37^. We analyze phonetic variation of the cardinal vowels in ten typologically diverse languages of the VC-C sample (Basque, Bulgarian, Hindi, Italian, Portuguese, Romanian, Russian, Tamil, Turkish, Vietnamese) and compare it with the corresponding variation in the MP-E sample. First, we pool the data of the 10 VC-C languages to evaluate within-language variation in the sample (see Methods). Figure 4 shows that the pattern of within-language variation in the VC-C sample (grey ellipses) is similar to that in the MPE sample (dashed grey ellipses) (multivariate comparison of variances: *p*=0.64; see Methods). Since the MP-E phonetic variation is conditioned by vocal tract variation (Fig. 3A), we infer that there exists a common mode of phonetic variation within languages that ultimately reflects a common mode of vocal tract variation. Second, we use the centroids from the 10 languages to evaluate the pattern of between-language variation in the VC-C sample (Fig. 4, red ellipses). Together, these analyses show that variation between languages is similar to variation within languages (red and grey ellipses in Fig. 4, respectively) (multivariate comparison of variances: *p*=0.36), suggesting that similar processes govern the phonetic diversification within and between languages. < Insert Figure 4 about here >

**Figure 4.**
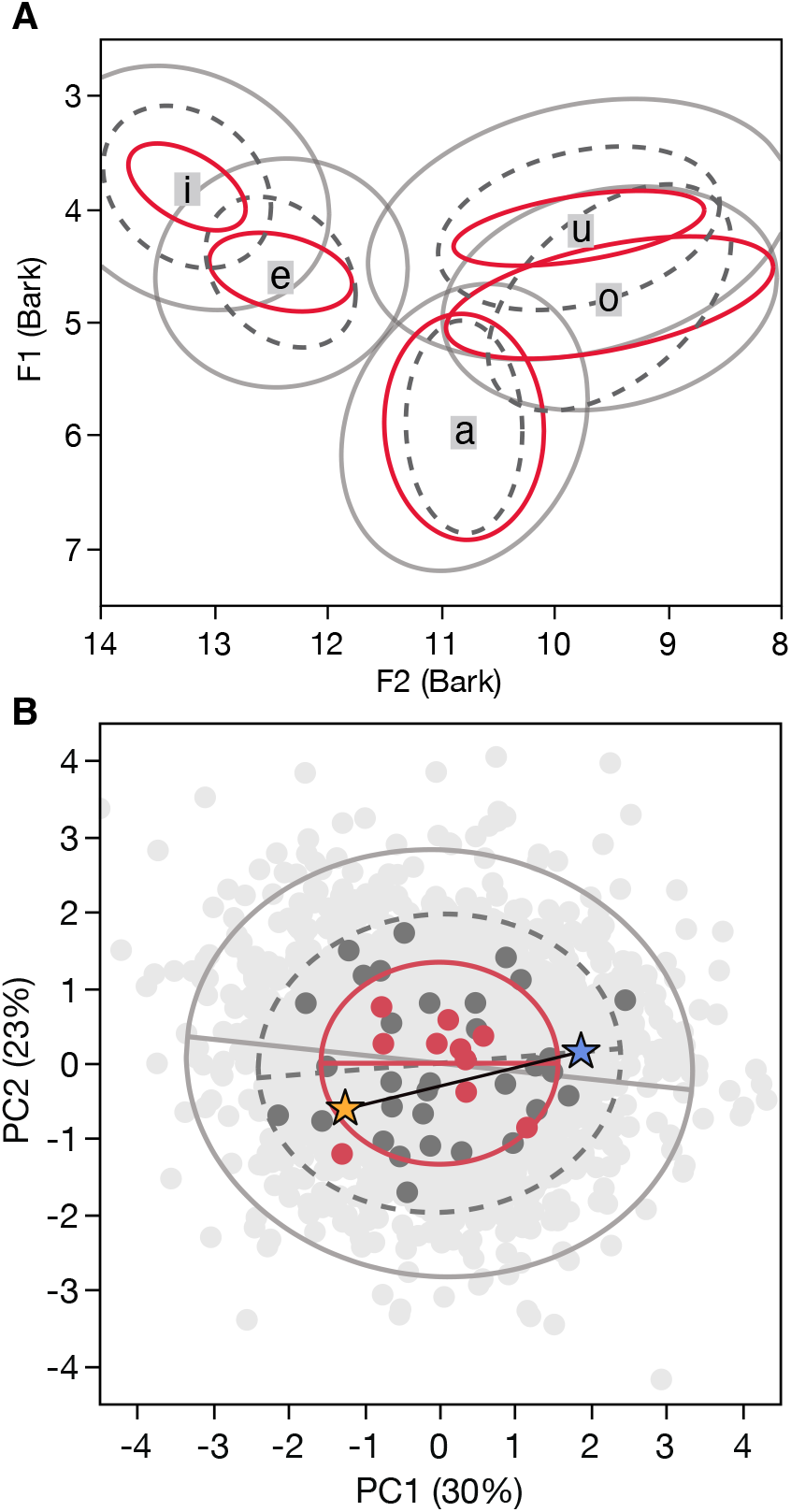
Patterns of within-language and between-language variation in cross-linguistic speech acoustics data. **A**: Acoustic variation of the cardinal vowels in F1-F2 formant space. 90% density ellipses show variation for the morpho-phonetic English speaker data (MP-E) (dark grey dashed lines), the pooled within-language data from the VoxCommunis Corpus (VC-C) (light grey solid lines) and the between-language VC-C data (red). Note the similarity in direction of variation across the different groups, which indicates a common mode of variation across the different data subsets. **B**: PCA of vowel polygon variation (90% density ellipses with major axes; colors as in **A**). Grey dots represent individuals (light grey: VC-C, dark grey: MP-E); red dots represent VC-C language means. The line connecting the yellow and blue stars marks phonetic variation due to vocal tract variation (MP-E data from Fig. 3A).

## Discussion

Many studies have taken a top-down approach to investigate language diversification, using methods of phylogenetic inference to reconstruct language family trees^38–40^. As a complement, we take a bottom-up approach to track how individual vocal tract variation in a speaker group serves as a source of articulatory and phonetic variation, and how phonetic variation within groups serves as a source of phonetic diversification between groups. At the individual level, we found that articulatory gestures and corresponding acoustics have similar vowel polygon representations in morphometric and phonetic spaces, respectively. While articulatory vowel polygons have not yet been reported, broadly similar vowel polygon representations have been found in neuromotor and neurosensory analyses^41–49^. Further studies are needed to elucidate the complete sequence of vowel representations, from neuromotor pattern generation to articulatory movements, acoustic output and speech perception^50, 51^.

We have also shown that the morphology of an individual’s vocal tract leaves a consistent signature on their articulatory gestures and the associated phonetic output. Previous studies have shown that speech sounds convey information about the physical, psychological, and social characteristics of the speaker^17, 52^. The morpho-phonetic signature described here is independent of these external variables and represents a common pattern of inter-speaker phonetic variability. This latter type of phonetic variability among speakers, however, is typically regarded as “noise” that hampers categorization tasks^53^ (e.g., automatic speech recognition) as well as phyletic reconstruction^54^. We argue that this so-called noise is an important source of phonetic variation between speaker communities, recalling a central tenet of evolutionary biology: within-group variation feeds processes of between-group diversification. Specifically, populations diversifying under a neutral (random) evolutionary process are expected to exhibit similar patterns of within- and between-population variation^55, 56^. In our case, variation in vowel phonetics within (*V*_pho.w_) and between (*V*_pho.b_) languages are indeed proportional to each other (Fig. 4A and Supplementary Fig. 9; see Materials and Methods). Furthermore, we showed that within-group phonetic variation is proportional to vocal tract variation (Fig. 3). Together, our results support the hypothesis that the morpho-phonetic covariation revealed in Fig. 3 is a universal biological constraint that not only shapes the phonetic diversity within speaker communities, but also governs a process of neutral-like phonetic diversification between languages (Fig. 4).

The neutral-like vowel evolution (NVE) model that we propose here is meant to serve as a null hypothesis that helps to disentangle the biological versus cultural mechanisms governing the evolution of spoken languages^57^. While there is consensus that phonetic and phonological change within and diversification among languages is largely driven by socio-cultural factors^58^, empirical evidence has led to the hypothesis that changes in vocal tract morphology (be it through genetic, environmental and/or cultural factors) can change the phonological repertoires of languages^13, 59–61^. Model considerations propose that small vocal tract differences between speaker groups result in small phonetic differences (so-called “weak biases”), which tend to become amplified through transgenerational learning^21^. Weak-bias amplification (WBA) models typically assume that all individuals of a speaker group have the same vocal tract morphology^21^. However, our empirical data do not support this assumption (Figs. 1, 3 and Supplementary Figs. 4–8). Thus, WBA models cannot explain intergroup phonetic diversification under conditions of high intragroup variability.

The NVE model starts with the observation that vocal tract and phonetic variation is already present within every population and even exceeds between-population variation (Fig. 4). Moreover, the NVE model neither requires nor excludes vocal tract and/or environmental change as a condition of phonetic diversification between languages. Within the envelope of morpho-phonetic covariation, language communities are free to travel through phonetic vowel space (Fig. 4) in a random-like neutral manner, while deviations from neutrality are indicative of non-random phonetic changes induced by environmental and/or cultural factors.

It is often assumed that the biological “language evolution” was completed with the acquisition of the language faculty, and that all subsequent “language change” is due to socio-cultural processes^58^. This view has recently been challenged on both methodological and empirical grounds, indicating that biological processes still influence the course of spoken language evolution^13, 21, 62^. The present study adds to this evidence by showing that a key mechanism of biological evolution – within-group variation generating between-group diversity – is still at work in the phonetic diversification of languages.

## Acknowledgements

SM and TD were funded by the Swiss National Science Foundation (PCEFP1_186841, EVOPHON).

## Author contributions

All authors designed the study, interpreted the results, and wrote the paper. TD and SM conducted the analyses. TD and CZ designed the figures.

## Materials and Methods

### Morpho-phonetic data of English speakers (MP-E)

We used a subset of the USC 75-Speaker Speech MRI Database^33^, which consists of video and concurrent audio recordings from Real-Time Magnetic Resonance Imaging (rtMRI) of the upper vocal tract from an open-source database of linguistically motivated speech tasks. The rtMRI data were acquired at 83.28 frames per second at an image matrix size of 84×84 pixels. Synchronous audio recordings were obtained in the scanner using fiber-optic microphones and then were subsequently denoised and are available in .wav format.

We extracted carrier words (‘beat’, ‘bait’, ‘pot’, ‘boat’, ‘boot’) to analyze American English vowels (i,e,a,o,u) as produced (each twice) by *N*=32 speakers (16 females and 16 males; aged 18–59y) from different ethnic and linguistic backgrounds. For each speaker, MRI frames were extracted from the rtMRI videos (see Supplementary Fig. 1). Depending on the length of the vowel, around 10 consecutive images (corresponding to 0.12 seconds) were extracted. In total, 3210 rtMRI images were used for the morphometric analyses (32 speakers, ∼100 images each across two repetitions of five vowels). The audio data were synchronized with the extracted image positions for acoustic phonetic analysis of the vowels. Thus, each extracted set of images and synchronous timestamps in the audio recordings together represent a vowel visually and acoustically.

### Geometric morphometric analysis of vocal tract shape

The rtMRI images were resampled at 164×164 pixels for geometric morphometric (GM) analysis^32, 63^. The shape of the upper vocal tract was quantified with *K*=95 two-dimensional landmarks, denoting both stationary skeletal and soft tissue structures (philtrum, anterior nasal spine, upper incisor tip, hard palate, inferior end of the nasopharynx, anterior borders of intervertebral disks between vertebrae C2 to C5) and moving skeletal and soft-tissue structures (lips, chin, supramentale, tongue, soft palate, uvula, epiglottis). The landmarks consisted of anatomical reference points (*K*_*f*_=21) and semilandmarks (points along curves between reference points; *K*_*c*_=74). See Supplementary Fig. 2 and Supplementary Table 1.

Landmark positions were digitized semi-automatically on each of the 3210 rtMRI images using the R software packages *StereoMorph, Morpho* and *geomorph*^64–67^. Generalized Procrustes Analysis (GPA)^63^ was carried out to scale individual landmark configurations to unit centroid size and to minimize differences due to rotation and translation of the individual configurations. GPA was performed only with those hard-tissue landmarks that remain stationary during articulation (see Supplementary Fig. 2 and Supplementary Table 1). For each vowel of each speaker, a mean configuration was calculated. The morphometric data matrix used for all further analyses thus consisted of 5*N*=160 rows (representing the 5 vowels per *N*=32 individuals) and 2*K*=190 columns (representing the 2D Procrustes shape coordinates of the *K*=95 landmarks per configuration). Principal Components Analysis (PCA) was then performed on the Procrustes shape coordinates to create a low-dimensional representation of shape space. One major advantage of GM analysis is that each articulatory shape of the vocal tract (per individual and per vowel) is represented by one multidimensional point in shape space. Conversely, each point in shape space can be re-transformed into its corresponding vocal tract shape in 2D physical space. This property facilitates the identification and visualization of major patterns of shape variation in the sample (see Figs. 1–3).

### Acoustic data extraction

Vocal tract resonance frequencies (formants F1–F4) and the fundamental frequency (F0) were extracted from the audio recordings of the rtMRI videos at synchronous timestamps with the image frames. Vowel measurements were taken through visual inspection of the spectrograms using Praat^68^. This procedure is common practice given the unreliability of automatic formant trackers^69^, particularly in noisy environments (e.g., MRI scanner noise is present in higher frequency ranges).

An average length of 0.12–0.15 seconds of audio (corresponding to 10–12 video frames) was analyzed for each vowel. Vowel mid-points in utterances were used to reduce possible formant transition effects due to coarticulation. For the diphthongs ‘bait’ and ‘boat’, we extracted the formants of the first vowel (e, o) at its midpoint. Definitive vowel identification was cross-validated with the articulatory patterns in the rtMRI image frames at concurrent time stamps. Mean fundamental frequency (F0) was measured for every speaker and every vowel similarly to formant extraction. The formant measurements from the two rounds of recordings were then used to calculate a mean formant value averaged over both recordings for each vowel and each speaker.

Lastly, given the noise emitted by the MRI scanner, the acoustics of the speech recordings may contain features of the Lombard effect^70, 71^. However, since the articulatory and acoustic data are recorded simultaneously, it is reasonable to assume that both data sets are similarly biased, such that morpho-phonetic covariation remains unbiased. Nevertheless, future studies may benefit by improving the sound quality of intra-MRI audio recordings towards more natural settings.

### Morpho-phonetic co-analysis

#### Data preparation

Sex and vocal tract size are known to be important factors influencing the speech signal^23, 27, 72, 73^. Indeed, preliminary analyses of our morpho-phonetic data showed significant effects of sex and vocal tract size on vocal tract shape and acoustic output. In this study, we focus on those patterns of interindividual articulatory and acoustic variation that are independent of sex and vocal tract size and are thus common to all individuals in the sample. To remove the effects of sex and size from the data, we proceeded as follows. The vocal tract shape data (PCs 1-10) as well as the acoustic data (formants F1-F4 and fundamental frequency F0) were regressed against centroid size and sex. The respective technique (multivariate multiple regression) is analogous to a multivariate analysis of covariance (MANCOVA) and was performed with the R packages *geomorph* and *RRPP*^67, 74^. All subsequent analyses were then performed on the residuals of the regression data.

#### Correlations between articulatory and phonetic distances

Correlations between articulatory and phonetic distances (Fig. 1F) were evaluated by regressing, for each individual, the *N*=10 Euclidean distances between the five vowels in acoustic space versus articulatory space. To compute the distances in acoustic space, we used formant wavelengths rather than frequencies. This is based on the consideration that the vocal tract acts as a resonating body, such that articulatory changes to its dimensions tend to elicit proportional changes of formant wavelengths rather than frequencies.

#### Covariance between articulatory and acoustic data

Both in multidimensional articulatory and acoustic space, individual vowel polygons can be characterized by three properties: a) location in space, as given by the coordinates of the polygon centroid (i.e., the average position of all vowels), b) polygon size, quantified as its centroid size (the square root of the sum of the vowels’ squared distances from the centroid); and c) polygon shape (the vowel coordinates relative to the centroid location). These geometric properties serve as proxies for the following real-life morphological and acoustic properties: in morphological terms, polygon location is a proxy of the resting state morphology of the vocal tract (the average of all vowel articulations), polygon size corresponds to the articulatory range, and polygon shape characterizes the articulatory contrasts between vowels. In acoustic terms, polygon location corresponds to the timbre of the voice, size to the acoustic range, and shape to the acoustic contrasts between vowels. Partial Least-Squares (PLS) analysis was used to study the covariation between the articulatory and acoustic vowel polygon data, and to explore and visualize how individual vocal tract shape influences the acoustic output. PLS permits the exploration of the covariation between two blocks of multidimensional data. Here, the multidimensional data are *N* x 2*P* matrices of articulatory landmark data (*N*=number of speakers, 2*P*=the 2D coordinates of *P* landmarks), *N* x 5*Z* matrices of articulatory data (*N*=number of speakers, 5*Z*=number of vowels x number of shape PCs) or *N* x 5*Q* matrices of acoustic data (*N*=number of speakers, 5*Q*=number of vowels x number of formants). We used PLS to evaluate the covariation between palate shape (*P*_p_=31) and tongue shape (*P*_t_=50) (Fig. 2), and between the vowel polygons in articulatory space (5*Z*=50) and in acoustic space (5*Q*=10) (Fig. 3). Similar to PCA, PLS is a dimension reduction technique. It uses singular value decomposition of the covariance matrix of two data blocks (i.e. data matrices) to extract pairs of orthogonal components (termed here PLS component pairs) that capture the largest, second-largest, etc., proportions of the total covariance in the sample. Due to the high dimensionality of the data, significance levels for PLS covariance (*p*_pls_) were estimated empirically with the permutation procedures (1000 iterations) implemented in the R package *geomorph*^67^. We also evaluated the covariance ratio (CR)^75^, which is a dimension-independent measure of the degree of association between two sets of multivariate data. The CR quantifies the covariance between two multidimensional data blocks as a fraction of the covariance within each block.

#### Data visualization and rendering

To visualize the correlated articulatory-acoustic (morpho-phonetic) variation, we proceeded as follows. For Figure 2, we evaluated the average tongue and palate shape of the three most extreme individuals at opposite ends of the data scatter (Fig. 2A). To generate the vocal tract profiles (Fig. 2C) of the two extremes, we calculated the distances between 21 tongue landmarks (tongue tip to tongue back) and the palate outline. Distances were measured perpendicular to the central axis of the air tube using ImageJ^76^. A similar approach was used for Figure 3, where the average vocal tract and formant configurations of each three individuals at the opposite ends of the data scatter were calculated (Fig. 3A), and visualized in Figs. 3A–C.

### Cross-linguistic phonetic data (VC-C)

In order to compare the patterns of group-level morpho-phonetic variation in the MP-E sample with patterns of language-level variation, we used the VoxCommunis corpus^37^ (VC-C), a large-scale cross-linguistic data set, which builds on a massively-multilingual collection of transcribed speech from the publicly available Mozilla Common Voice Corpus^77^. The VC-C dataset includes acoustic-phonetic measurements (F1-F4) of 36 different languages. From these, we selected the languages that included the five cardinal vowels (i,e,a,o,u) that we used in our morpho-phonetic analyses of the MP-E data. We excluded data with obvious errors or too few speakers. We analyzed data points only from speakers who produced the full set of vowels (i,e,a,o,u), in order to get the full vowel polygon of each speaker. This led to a sample of 19,335 data points (vowel utterances), comprised of 3867 individuals from 10 different languages (Basque, Bulgarian, Hindi, Italian, Portuguese, Romanian, Russian, Tamil, Turkish, Vietnamese). To directly compare the VC-C data with our MP-E data, we adjusted the data for differences in formant extraction methods by superimposing vowel centroids. Furthermore, we ran similar data adjustment procedures as for the MP-E in order to obtain sex-adjusted between-language data (pooled for sex, *N*=19,335). Variation in the sex-pooled phonetic data was then decomposed into variation within languages (*V*_pho.w_), i.e., shared by all languages and variation between languages (*V*_pho.b_). *V*_pho.w_ was evaluated by pooling the data for language (*N*=19,335), while *V*_pho.b_ was evaluated from the mean data per language.

### Comparing patterns of variation

#### Graphical comparison

We compared the principal patterns of phonetic variation in the MP-E sample and in the VC-C sample (both within-languages and between-languages) by projecting all data into formant space (Fig. 4A; variation per vowel) as well as into a common PC space (Fig. 4B; variation of the vowel polygon). To construct the graph of Fig. 4B, the vowel polygon data were expressed as data vectors of length 5*Q* (5 vowels, *Q*=2 formants). PCA was performed on the 10×10 data matrix of the mean vowel polygons of the 10 languages, such that PC1 of Fig. 4B expresses the principal direction of between-language variation of the vowel polygon. The principal directions of within-language variation in the VC-C and MP-E samples were projected into the same PC space. Finally, the principal direction of morpho-phonetic covariation in the MP-E sample (acoustic PLS1 in Fig. 3A) was also projected into the PC space.

#### Statistical comparison of variance-covariance structure

The near-collinearity of the principal variation vectors in Fig. 4B indicates proportionality of the variance-covariance matrices underlying these distributions. Statistical tests on proportionality were performed with eigenanalysis^78^, as implemented in the R package *vcvcomp*^79^ (see *Vocal tract variation influences within- and between-language phonetic diversity* in main text).

*Test for neutral vowel evolution*. In analogy to the standard model of neutral phenotypic evolution, we expect that, under neutral conditions of language change, the pattern of vowel polygon variation between languages is proportional to the variation within languages, *V*_phon.b_∼ *V*_phon.w_^55^. This proposition can be tested as follows^56^: the VCV (variance-covariance) matrices characterizing between-language and within-language variation are each orthogonalized, yielding statistically independent variance components along the eigenvectors. Then the between-group variance components are regressed against the within-group components. Proportionality between the two VCV matrices is indicated by linearity (i.e., slope = 1 in a double-logarithmic plot; Supplementary Fig. 9).

## Supplementary Figures

**Supplementary Figure 1.**
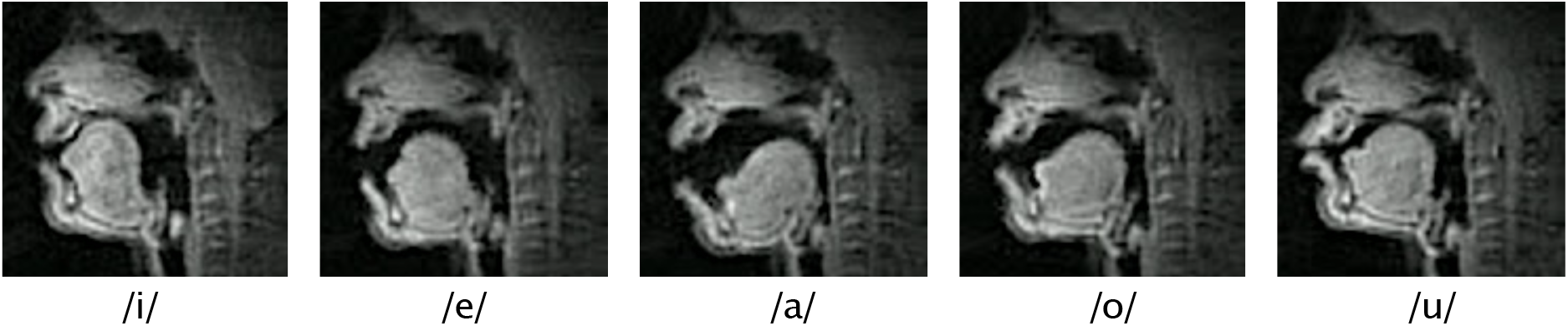
Example of rtMRI image frames during vowel production.

**Supplementary Figure 2.**
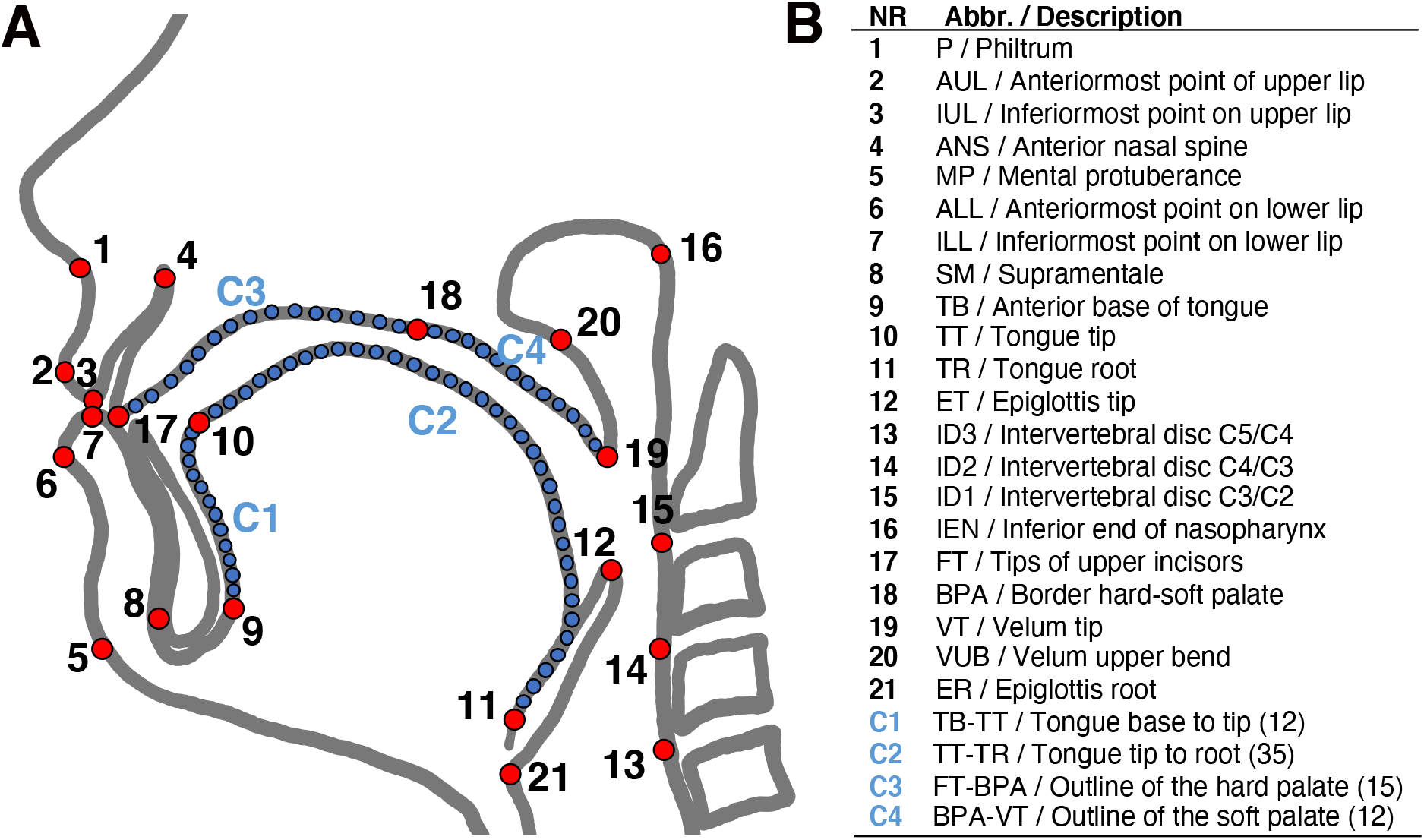
Schematic representation of landmarks placed on 2D midsagittal images of the vocal tract. **A**: Anatomical landmarks (reference points) are in red and semilandmarks in blue. **B**: Landmark definitions. See Supplementary Table 1 for detailed descriptions of the location of the landmarks.

**Supplementary Figure 3.**
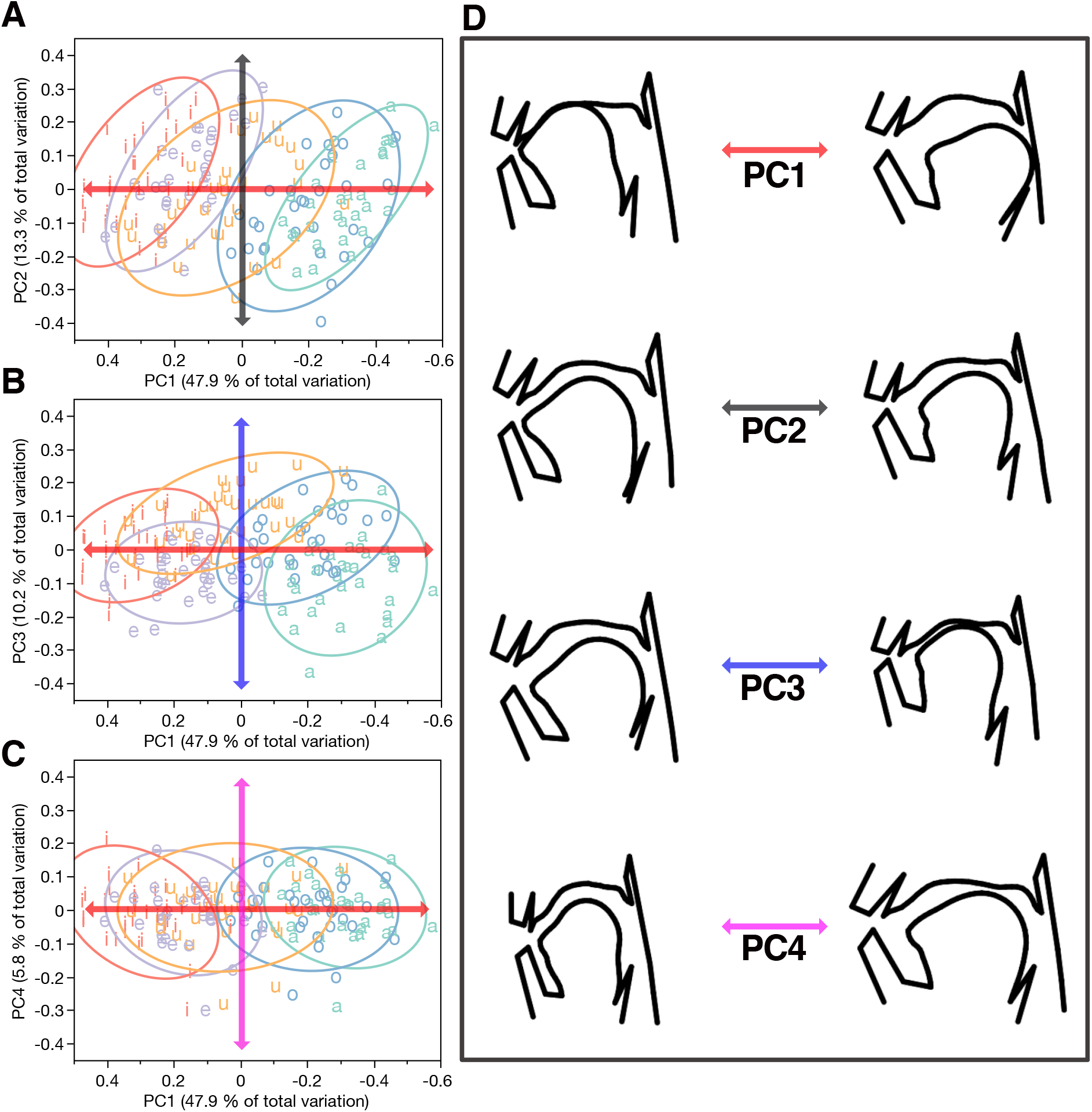
Articulatory shape variation associated with the PCs of shape space. **A**: Shape variation along PCs 1 and 2. **B**: Shape variation along PCs 1 and 3. **C**: Shape variation along PCs 1 and 4. **D**: Visualization of vocal tract shape variation along PCs 1-4 at the two ends of the respective data scatter.

**Supplementary Figure 4.**
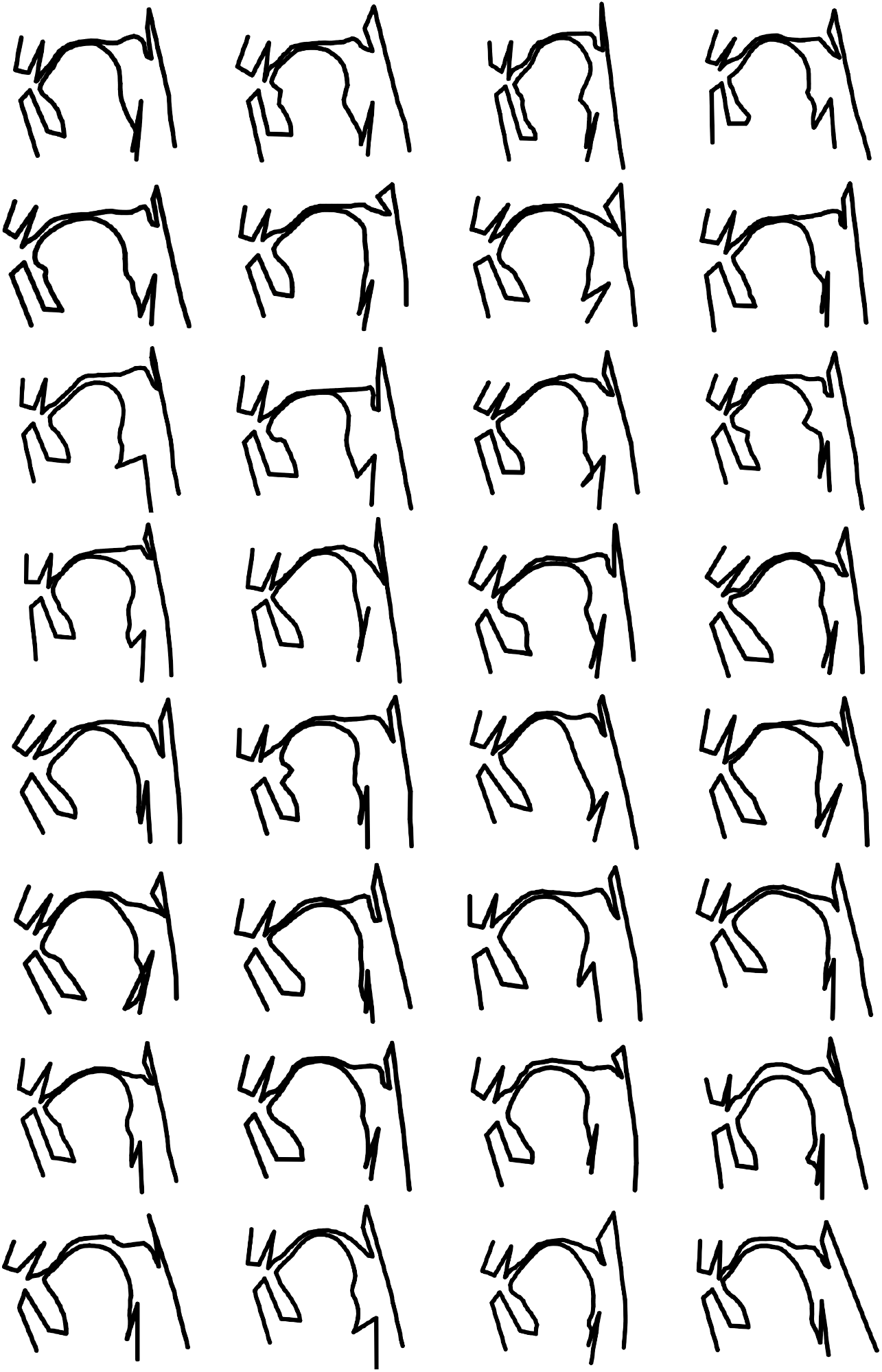
Physical shape variation in the full sample of speakers producing the vowel [i].

**Supplementary Figure 5.**
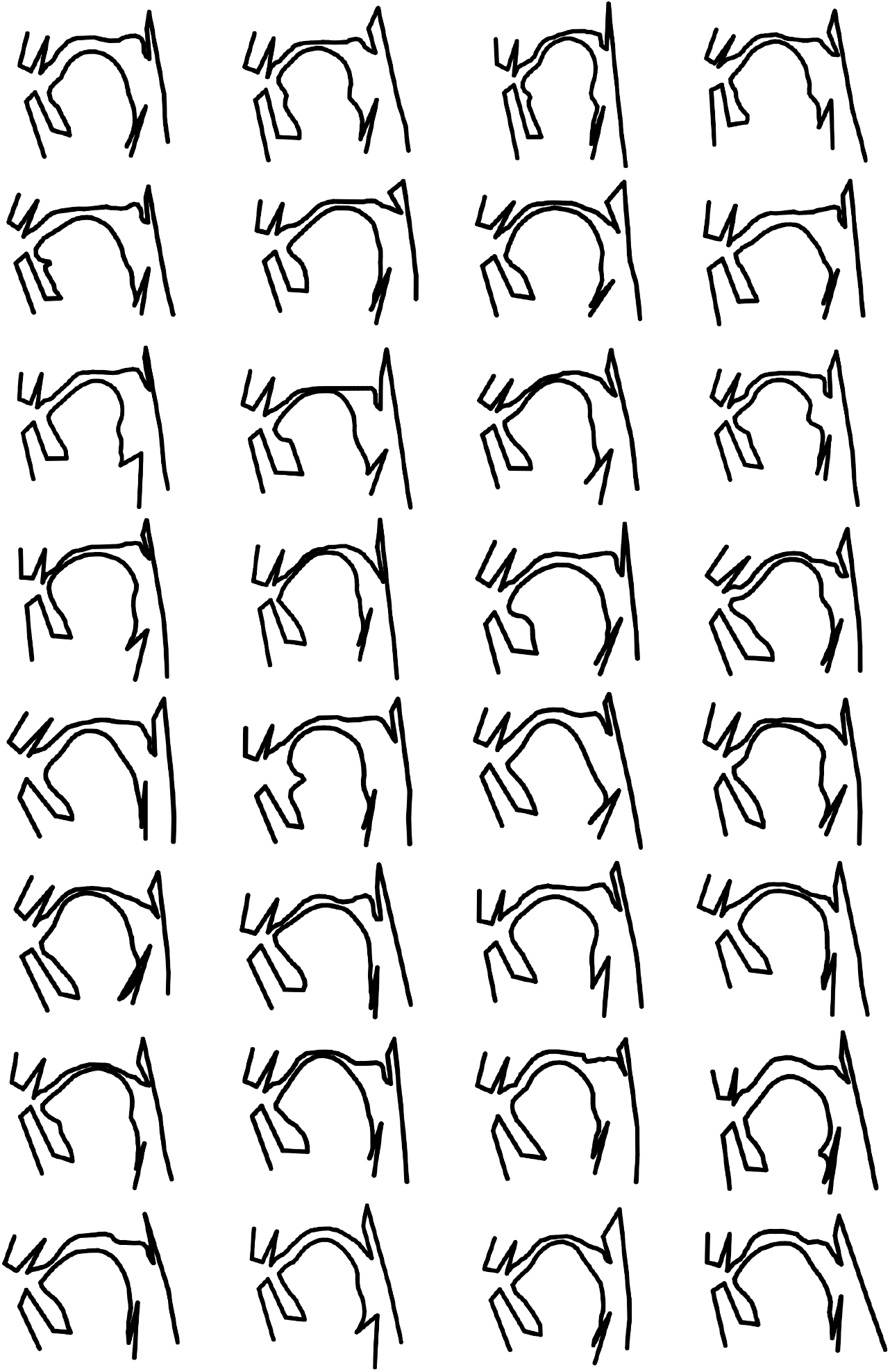
Physical shape variation in the full sample of speakers producing the vowel [e].

**Supplementary Figure 6.**
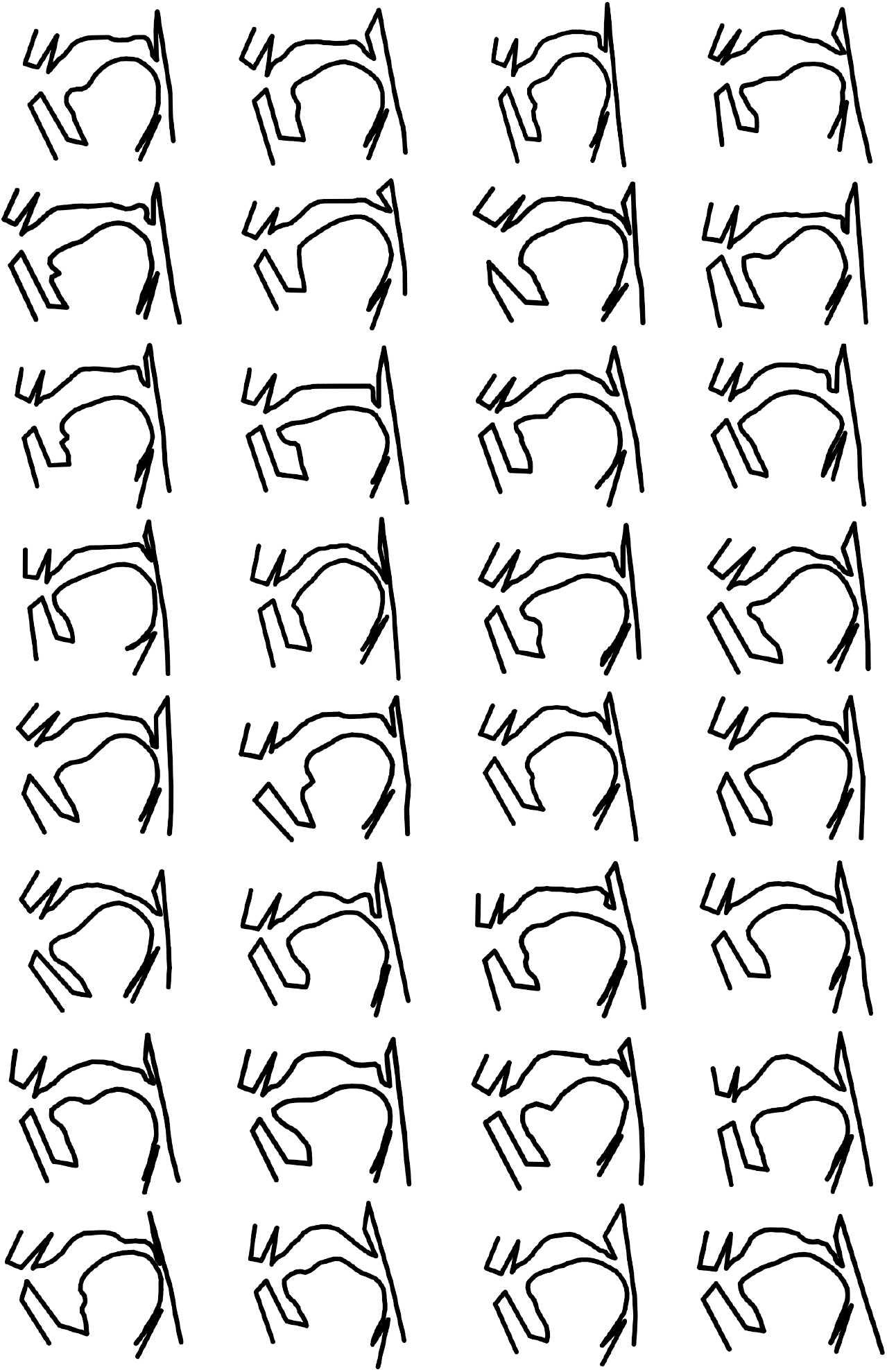
Physical shape variation in the full sample of speakers producing the vowel [a].

**Supplementary Figure 7.**
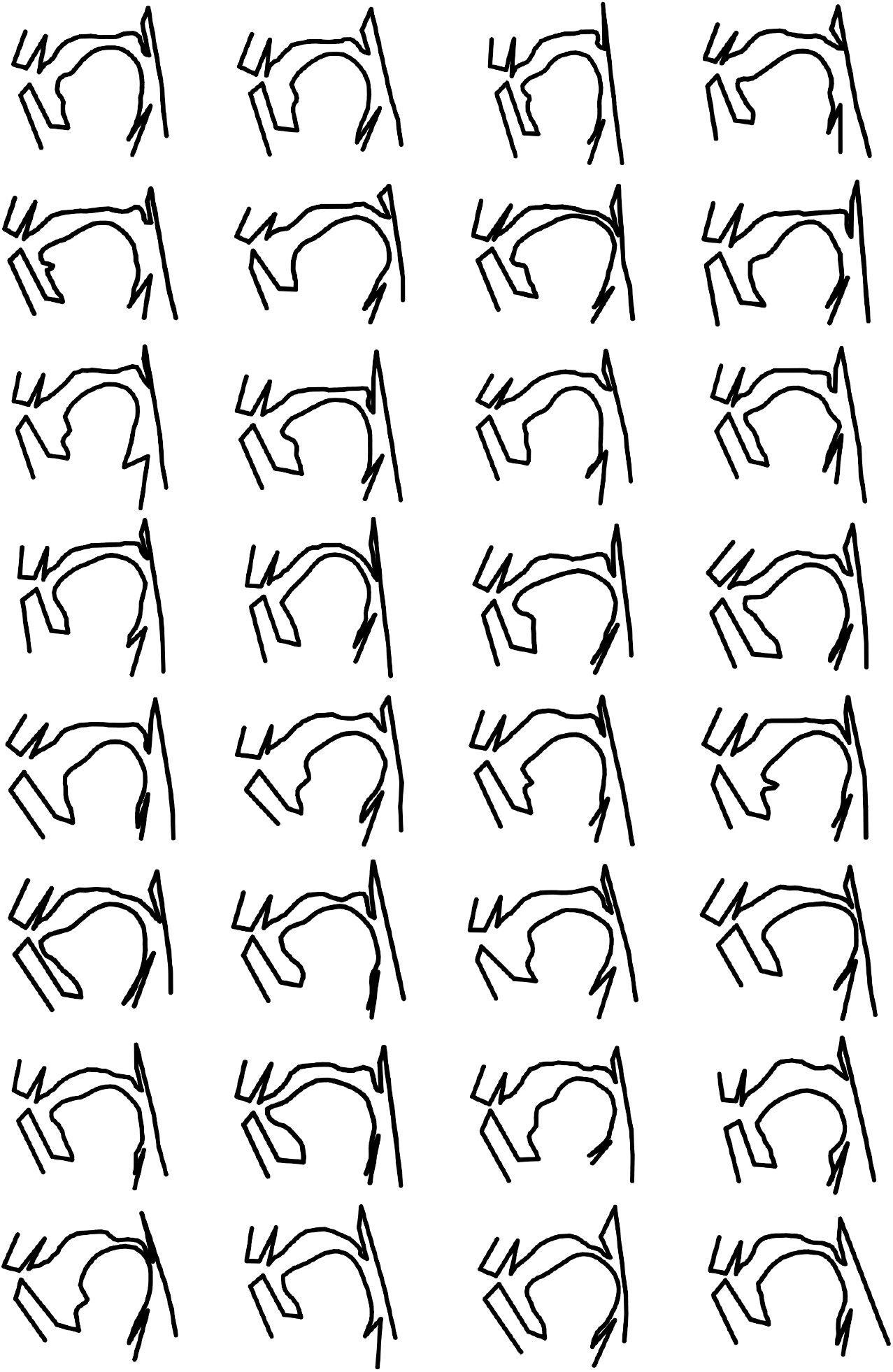
Physical shape variation in the full sample of speakers producing the vowel [o].

**Supplementary Figure 8.**
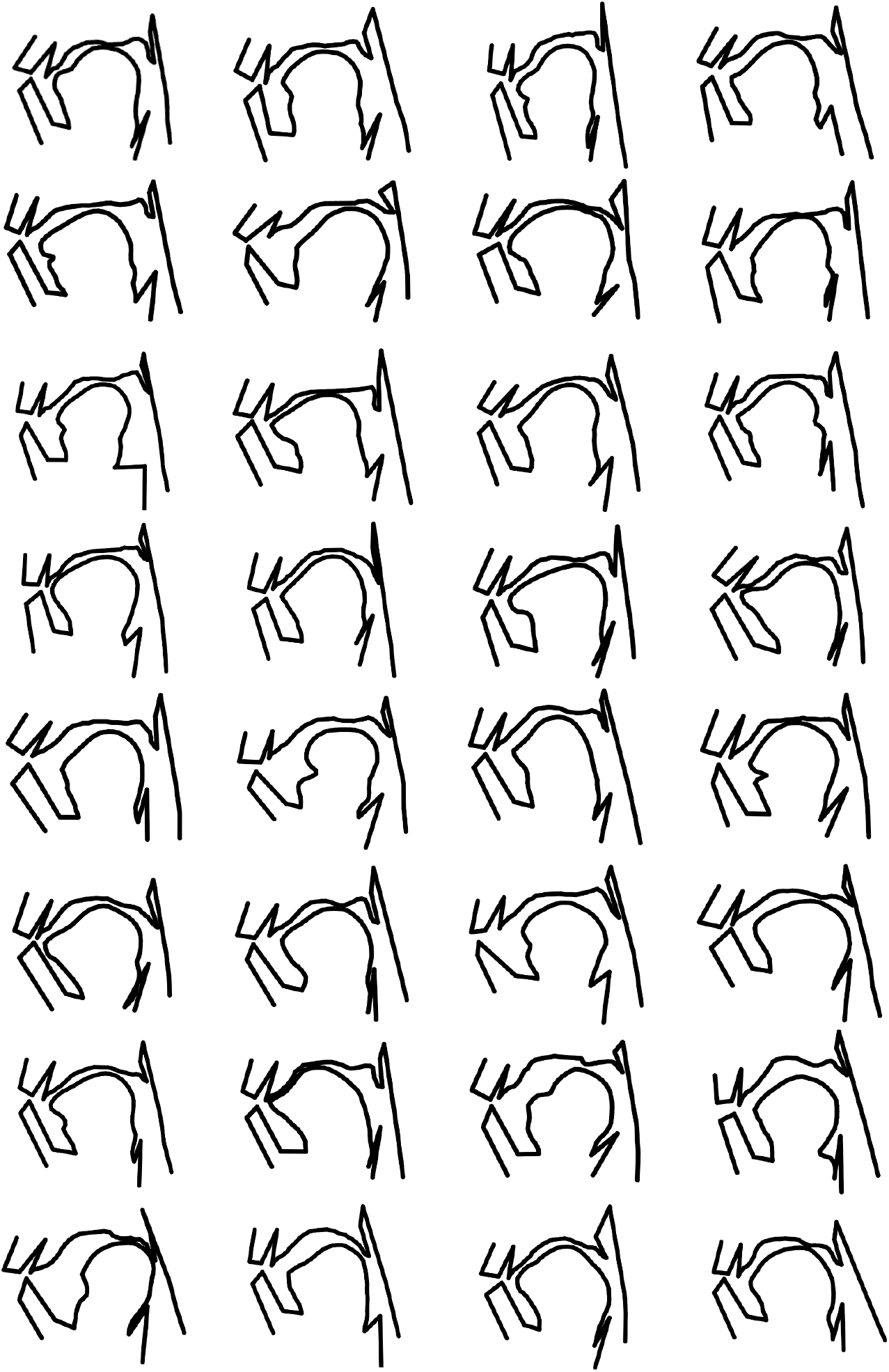
Physical shape variation in the full sample of speakers producing the vowel [u].

**Supplementary Figure 9.**
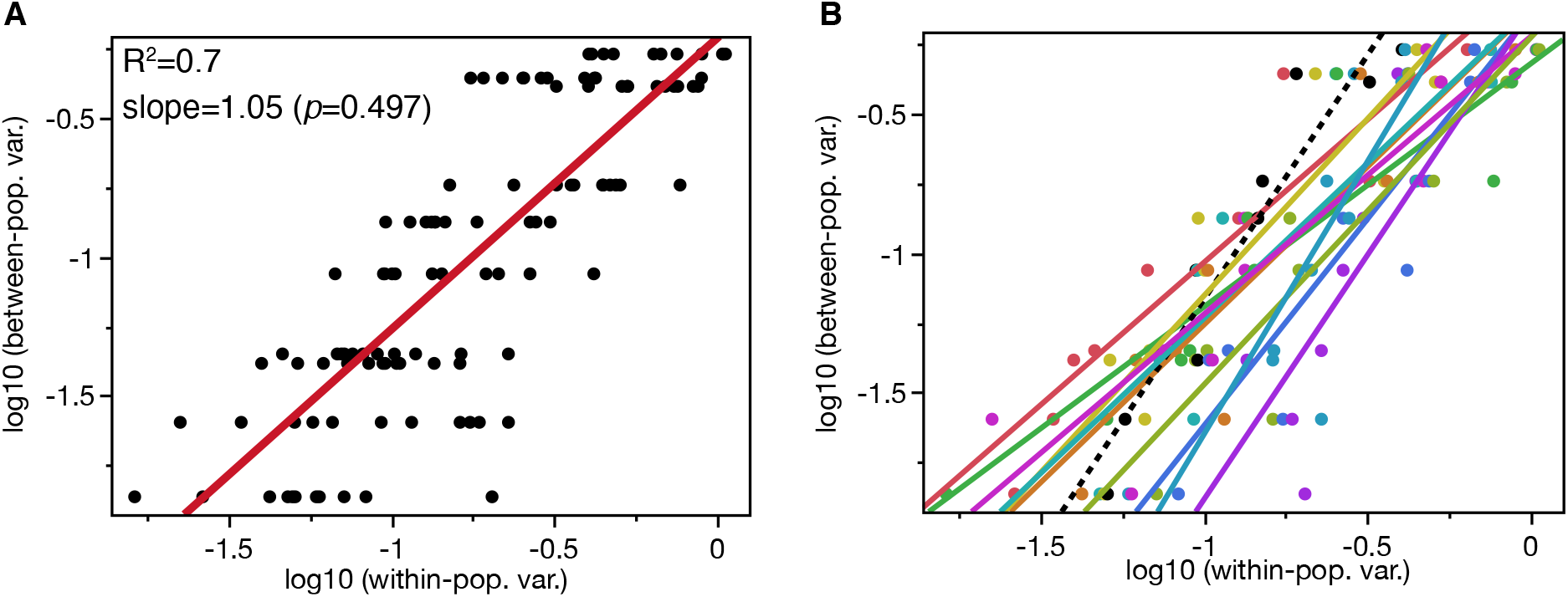
Regression of between-population (between-language) variances on within-population variances. **A**: Regression of between-language variance on within-population variance by means of PCs 1-10 of vowel polygon form in F1-F2 formant space. The slope does not deviate significantly from linearity (*p=*0.497) The within-population variances were computed on the MP-E data and the VC-C data. **B**: Separate regression for each VC-C language (each color corresponds to a different language) and the MP-E (black dashed line).

## Supplementary Tables

**Supplementary Table 1.** Orofacial landmarks (LM) used in this study.

| ID | Code | Description | K | Type | Tissue |
| --- | --- | --- | --- | --- | --- |
| 1 | P | Philtrum | 1 | F | H |
| 2 | AUL | Anteriormost point on upper lip | 1 | F | S |
| 3 | IUL | Inferiormost point on upper lip | 1 | F | S |
| 4 | ANS | Anterior nasal spine | 1 | F | H |
| 5 | MP | Mental protuberance | 1 | F | H |
| 6 | ALL | Anteriormost point on lower lip | 1 | F | S |
| 7 | ILL | Inferiormost point on lower lip | 1 | F | S |
| 8 | SM | Supramentale | 1 | F | H |
| 9 | TB | Anterior base of tongue | 1 | F | S |
| 10 | TT | Tongue tip | 1 | F | S |
| 11 | TR | Tongue root | 1 | F | S |
| 12 | ET | Epiglottis tip | 1 | F | S |
| 13 | ID3 | Intervertebral disc C5/C4 | 1 | F | H |
| 14 | ID2 | Intervertebral disc C4/C3 | 1 | F | H |
| 15 | ID1 | Intervertebral disc C3/C2 | 1 | F | H |
| 16 | IEN | Inferior end of nasopharynx | 1 | F | H |
| 17 | FT | Tips of upper incisors | 1 | F | H |
| 18 | BPA | Border hard-soft palate | 1 | F | S/H |
| 19 | VT | Velum tip | 1 | F | S |
| 20 | VUB | Velum upper bend | 1 | F | S |
| 21 | ER | Epiglottis root | 1 | F | S |
| Curve1 | TB - TT | Tongue base to tip | 12 | C | S |
| Curve2 | TT - TR | Tongue tip to root | 35 | C | S |
| Curve3 | FT - BPA | Outline of the hard palate | 15 | C | H |
| Curve2 | BPA - VT | Outline of the soft palate | 12 | C | S |

**Supplementary Table 2.** Variance ratios of within-vowel variance vs. between-vowel variance.

| Vowel | VR<sup>Articulatory</sup> | VR<sup>Formant</sup> |
| --- | --- | --- |
| **a** | 0.546 | 0.064 |
| **e** | 0.501 | 0.040 |
| **i** | 0.495 | 0.053 |
| **o** | 0.585 | 0.087 |
| **u** | 0.611 | 0.079 |
| Mean | **0.548** | **0.065** |

